# Assessment of antimicrobial potential of cell-free supernatant from *Lacticaseibacillus paracasei* SRG isolated from the vaginal microbiota of healthy women against uropathogens

**DOI:** 10.64898/2026.10.11.758281

**Authors:** Riya Anand, Deepshikha Rai, Pradyot Prakash, Ganesh Kumar Maurya

## Abstract

Among women, urinary tract infection (UTI) represents a major health concern. Approximately 50% of the adult female population experienced an episode of UTI at least once. UTI and recurrent UTIs caused by resistant bacteria are very common in clinical settings. The treatment and management of UTI is becoming more challenging because of the rise in antibiotic-resistant bacteria. *Lactobacillus* spp are the dominant bacterial genus in urinary and vaginal microbiota, which are also used as probiotics. In this study, *Lacticaseibacillus paracasei* SRG isolated from the vagina of healthy women were evaluated for its probiotic-associated properties and cell-free supernatant (CFS)-mediated antibacterial activity against eight uropathogens. The antibacterial activity of CFS was concentration- and pH-dependent, which was significantly evident at CFS/2 and CFS/4 concentrations and acidic pH values of 4 and 5. The antibacterial activity in CFS remains unaltered even after heat and proteinase K treatment, which suggests the role of heat- and protease-resistant metabolites beside the contributions from acidic conditions. *L. paracasei* SRG showed strong autoaggregation (85.32) and cell-surface hydrophobicity (55.60). In addition, it also exhibits co-aggregation with all indicator strains, suggesting substantial interaction with diverse pathogenic bacteria. These observations highlighted the *L. paracasei* SRG as a potential probiotic candidate developed from indigenous vaginal microbiota.

## 1. Introduction

Urinary tract infections (UTI) are the most prevalent infection among women than males. More than 50% of adult women experience UTI at least once in their life and more than 20% of patients experience at least two episodes of UTI occurring within six months or at least three episodes within a 12-month period, which is called as recurrent UTI (rUTI) [1]. Women are more susceptible to UTI due to the proximity of urethra to the rectum. The anatomical proximity of the anus and the urogenital tract in females, facilitate the colonization of opportunistic microorganisms such as *Escherichia coli, Klebsiella pneumoniae, Enterococcus faecalis, Staphylococcus saprophyticus, Proteus mirabilis, Gardnerella vaginalis* and *Candida albicans* [2]. Many studies show that the urinary and vaginal microbiomes are closely interconnected, particularly in women with urinary disorders [3, 4]. Vaginal colonization by uropathogens is considered as a key precursor for UTI pathogenesis because of shared niche between the urinary tract and vagina. A pivotal study by Yuko M. Komesu et al. strengthened evidence for the existence of a shared urogenital microbiome by demonstrating significant overlap and strong correlations in bacterial communities between catheterized urine and vaginal samples [5]. The study identified *Lactobacillus* as the dominant bacterial genus in urinary and vaginal niches, constituting approximately 53% and 64% of the microbiome, respectively. Recently, comparative study of urinary microbiome from healthy controls and UTIs patients identified the predominance of *L. crispatus* in healthy controls and *E. coli* with UTI patients [6]. In addition, several other bacterial genera, including *Gardnerella, Prevotella, Ureaplasma*, and *Escherichia*, showed strong correlations between urinary and vaginal samples. Collectively, these observations support the concept of shared urinary and vaginal microbiomes which is vital for maintaining urogenital health [5]. Furthermore, alterations in this microbial homeostasis may contribute to recurrent UTIs and highlight the potential of probiotics and postbiotics as a microbiome-based therapeutic strategies. Several studies showed that some of the vaginal *Lactobacillus* spp. can inhibit the growth and biofilm formation in uropathogens through antimicrobial substances, primarily by organic acids [7]. Generally, the organic acids found in the cell free supernatant (CFS) after the fermentation of broth include lactic acid, acetic acid, citric acid, butyric acid and propionic acid by the *L. plantarum* and *L. crispatus* strains [8-10].

In the present study, we investigated the antimicrobial potential of the CFS of *Lacticaseibacillus paracasei* SRG strain isolated from healthy vaginal microbiota against clinically relevant uropathogens. We aim to study, whether metabolites produced by a naturally occurring vaginal *Lactobacillus* can directly inhibit uropathogenic bacteria, and to further characterise its antimicrobial compounds as potential non-antibiotic strategies for UTI prevention or management.

## 2. Materials and methods

### 2.1 Uropathogenic strains used and growth conditions

Eight uropathogenic strains were used, including five Gram-negative strains and three Gram-positive strains. The Gram-negative strains, *E. coli* BW25113, *E. coli* CRE (carbapenem-resistant clinical strain), *K. pneumoniae* KCTC12385, *K. pneumoniae* CRK (carbapenem-resistant clinical isolate), and *P. mirabilis* MCC4207 were used. The Gram-positive strains, vancomycin-resistant *E. faecium* ATCC700221 (VRE), methicillin-resistant *S. aureus* JE2 (MRSA) and methicillin-sensitive *S. aureus* ATCC25923 (MSSA) strains were used in this study.

For the experiment, the strains of *E. coli, K. pneumoniae*, and *S. aureus* were streaked onto Luria-Bertani (LB) agar plates while strains of *P. mirabilis* and *E. faecium* were streaked onto Tryptic soy agar (TSA). All these strains were incubated at 37°C. Further, strains were grown in their respective media broth at 37°C and 180 rotation per minute (RPM) under aerobic conditions for overnight.

### 2.2 Isolation of lactobacilli from the vaginal swab of healthy women

Six Indian women (18-40 years old) having no symptoms of urogenital tract infection in the past one year were selected for this study. All the volunteers provided a written informed consent as per the guidance of ethics committee, and proper instructions were provided for the self-swabbing vaginal sample method. The samples were coded to ensure full anonymousness. The samples were immediately processed for the isolation of lactobacilli. Firstly, the swabs were suspended in 1 mL of sterile phosphate buffered saline (PBS), serially diluted and spread onto de Man, Rogosa and Sharpe (MRS) agar plates. The plates were incubated under anaerobic conditions using Anoxomat® III anaerobic jar system for 24 h at 37°C. Next, multiple colonies were picked using sterile toothpick and suspended individually in 1ml of MRS broth for further growth under anaerobic conditions. Bacterial cultures were centrifuged at 10,000 RPM for15 minutes at room temperature and supernatant were collected for initial screening of antimicrobial property against *E. coli* BW25113 and CRE. Isolates showing significant growth reduction of uropathogens were selected for further characterization. Further Gram-staining was done using the protocol of Gram’s Colour Staining Kit (CDH Fine Chemical, Lucknow) and observed under the light microscope. Also, the catalase test was done for further confirmation.

### 2.3 Genome isolation and diagnostic PCR analysis

For genome isolation, the cetyltrimethylammonium bromide (CTAB) method was used [11]. Anaerobically grown culture of the isolated strain was centrifuged, and the pellet was dissolved in 200 µL TE (Tris-EDTA) buffer; then 10 µL 50mg/mL lysozyme was added and incubated for 5-6 h at 37°C. Then 70 µL 10% sodium dodecyl sulphate and 5 µL 20 mg/mL proteinase K were added and incubated at 65°C for 10 min. Then, 100 µL 5 M sodium chloride and 100 µL pre-warmed 10% (w/v) CTAB/700 nM sodium chloride buffer were added and incubated at 65°C for 10 min. Further, chloroform/isoamyl alcohol (24:1, v/v) was added in equal volume, and centrifuged at 10000 RPM for 10 min. The upper aqueous supernatant was mixed with 0.6 volume of chilled isopropanol and incubated at 4°C for 30 min. Next, tubes were spun at 10000 RPM for 10 min to discard the supernatant. Further, pellet was washed with 1 mL 70% ethanol through centrifugation. Finally, pellets were air dried and suspended in 80 µL ultrapure water followed by storage at -20°C.

### 2.4 16S rRNA gene amplification, Sanger sequencing and phylogenetic analysis

For 16S rRNA gene sequencing, we used universal primers: 27F 5’AGAGTTTGATCCTGGCTCAG and 1492R 5’ GGTTACCTTGTTACGACTT. The PCR was performed using Q5 DNA polymerase (New England Biolab) at T_m_ (melting temperature) of 48.9°C. The PCR products were cleaned using QIAquick PCR Purification Kit (QIAGEN, Germany), and subjected to agarose gel electrophoresis for quality check (Figure 1C). Subsequently, purified PCR products were processed for Sanger sequencing of the 16S rRNA gene region using 27F and 1492R primer set. The FASTA files containing the both forward and reverse sequence reads after Sanger sequencing were subjected to NCBI nBLAST tool for the identification of vaginal isolate. Further, phylogenetic analysis of identified bacterial strains was performed against *L. fermentum* MBO003 (MT071603.1), *L. paracasei* 1989 (MT597791.1), *L. crispatus* JCM 7697 (AB911456.1), *L. gasseri* DMBCT6 (KM056281.1), *L. iners* 109878T (HE573916.1) and *L. jensenii* TMPC 90217 (OM265435.1) using MEGA11 software as described in [12,13].

**Figure 1.**
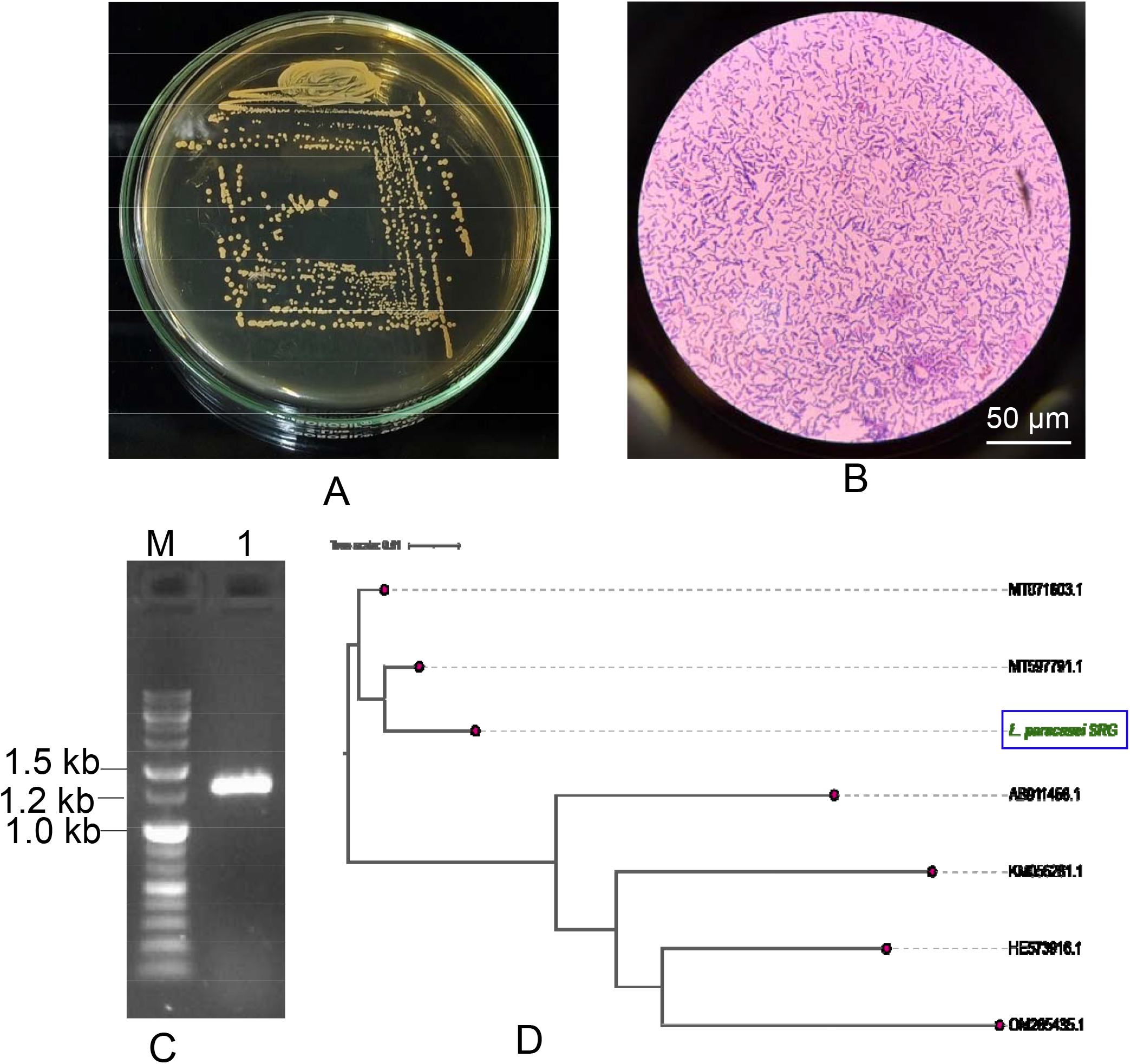
Isolation (A), Gram-staining (B), molecular identification using 16S rRNA gene Sanger sequencing (C) and phylogenetic positioning (D) of *L. paracasei* SRG isolated from the human vaginal swab. Lane M indicate DNA marker and Lane 1 shows the purified PCR product of 16S rRNA gene in figure (C).

### 2.5 Preparation and antimicrobial activity of CFS

The isolated vaginal *Lactobacillus* strain was cultured in MRS broth anaerobically for 48 h at 37°C. The grown culture was centrifuged at 8000 RPM at 37°C for 15 min and filtered through 0.22 µm filters to remove residual bacterial cells, if any. The antimicrobial activity of CFS against the selected uropathogenic bacterial strains was assessed in a 96-well plate as described in [10]. The supernatant was serially diluted in MRS broth to get four different serial dilutions with the CFS/MRS ratio of 0.5, 0.25, 0.12, and 0.06. Additionally, the pH of the supernatant was adjusted to different pH by adding different volumes of 1 N NaOH until four different conditions were achieved. The CFS pH includes, unaltered CFS pH 4, CFS at pH 5, CFS at pH 6 and CFS at pH 7. Furthermore, CFS was treated with proteinase K (20 µg/mL for 1h at 37°C) and heat (98°C for 1h) and analysed for its antimicrobial property against selected uropathogens. Different treatment conditions include CFS only, proteinase K treated CFS (CFS + Pr K), Heated CFS (CFS + Heat), Heated MRS (MRS + Heat), MRS pH4 and MRS only as untreated control. 100 µL of the different indicator strains with an initial colony forming units (CFU) of 5 × 10^7^ CFU/mL were incubated with 100 µL of the above-mentioned CFS conditions for 24 hours at 37°C and 150 RPM in a 96-well plate. Next, CFUs were measured after serial dilutions of all tested conditions across different uropathogenic indicator strains.

### 2.6 Assessment of autoaggregation, coaggregation and cell surface hydrophobicity

Autoaggregation, coaggregation, and cell surface hydrophobicity of the selected vaginal isolate (*L. paracasei* SRG) were evaluated as described earlier [14]. For autoaggregation and coaggregation assays, the vaginal isolate was cultured under anaerobic conditions in MRS broth for 48 h at 37°C, and centrifuged at 10000 RPM for 10 min at 37°C to collect cell pellet. Next, pellet was washed twice with PBS and resuspended in PBS. The bacterial suspension was adjusted to the optical density (O.D.) of 0.25 ± 0.05 at 600 nm. For autoaggregation assay, 1 mL aliquot of the suspension was incubated at 37°C without agitation. The OD□ □ □ was measured at 0 and 5 h to determine the extent of autoaggregation. The autoaggregation percentage was calculated using the following equation:

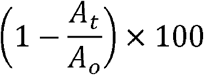

Where, ‘A_o_’ represents the initial absorbance at 600 nm and ‘A_t_’ represents the absorbance after 5 h of incubation.

For the coaggregation assay, selected uropathogens were also cultured in their respective media broth, harvested, washed and resuspended in PBS to adjust the O.D. of 0.25 ± 0.05 at 600 nm. Further, 1 mL of vaginal isolate and 1 mL of uropathogen at similar O.D. were mixed and incubated at 37°C without agitation for 5 h. The O.D. (at 600 nm) values of each bacterial suspension and their corresponding mixtures were recorded initially and end of the incubation period. The percentage of coaggregation was calculated using the following equation:

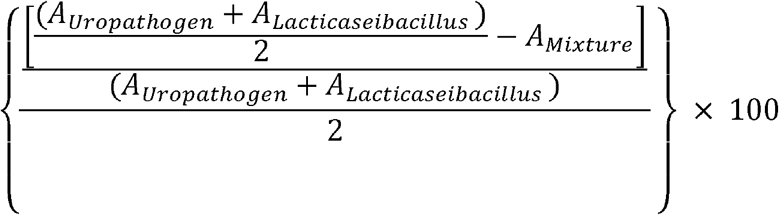

Where, ‘A_Uropathogen_’ and ‘A_*Lacticaseibacillus*_’ represent the absorbance values of the uropathogen and *Lacticaseibacillus* suspensions, respectively, and ‘A_Mixture_’ represents the absorbance of the mixed bacterial suspension after 5 h of incubation. The absorbance values of the individual suspensions at the corresponding time point were used for the calculation.

The cell surface hydrophobicity of the *L. paracasei* isolate was measured by microbial adhesion to hydrocarbons (MATH) method using xylene as the organic phase. The bacterial suspension was prepared as described above and adjusted to an OD 0.25 ± 0.05 at 600 nm. An equal volume of xylene was added, and the mixture was vortexed for 3 min. To facilitate phase separation, the mixture was incubated at room temperature for 1 h. Then the O.D. of the aqueous phase was recorded at 600 nm. Hydrophobicity was represented as the % adhesion to xylene and calculated as follow:

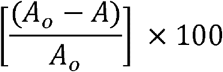

Where, ‘A_o_’ indicate the initial absorbance before adding xylene and ‘A’ is the absorbance of the aqueous phase after xylene extraction.

### 2.7 Statistical analysis

Data were presented as mean ± SD obtained from three independent replicates and analyzed by one-way ANOVA followed by Dunnett’s multiple comparison tests using GraphPad Prism software v8.0. Statistical significance was represented with the help of asterisks having following thresholds: *p* < 0.05 (*), *p* < 0.01 (**), *p* < 0.001 (***) and *p* < 0.0001 (****).

## 3. Results and discussions

### 3.1 Morphological and biochemical characterization

*Lactobacillus* is one of the important genera of group lactic acid bacteria which belongs to the phylum Firmicutes (Bacillota). Lactobacilli are non-spore-forming, catalase-negative, rod-shaped Gram-positive bacteria that majorly produce lactic acid as end metabolites of fermentation [10,15,16]. Microscopic analysis of the isolated strain shows rod-shaped, purple-coloured bacteria, confirming the strain is Gram-positive (Figure 1A,B). The isolate tested negative for catalase activity, whereas the control *E. coli* BW25113 strain exhibited a positive catalase reaction.

### 3.2 Molecular identification of vaginal isolate and phylogenetic analysis

The Sanger sequencing of amplified 16S rRNA gene identified the vaginal isolate as *Lactocaseibacillus paracasei* with a percentage identity of 98.69% in nBLAST search. We named this strain as “*Lactocaseibacillus paracasei* SRG”. The comparative phylogenetic tree reveals the relationship between the vaginal isolate ‘*L. paracasei* SRG’ and the representative *Lactobacillus* of different community strain type (CST) based on the 16S rRNA gene sequences. The phylogenic analysis suggested that the selected vaginal isolate belonged to the genus *Lactobacillus* and mainly to the phylogenetic clade of *L. paracasei* (Figure 1D).

### 3.3 Effect of *L. paracasei* SRG CFS on selected uropathogens

The analysis of antibacterial activity of CFS at different concentrations against selected uropathogens showed substantial reduction of bacterial growth (Figure 2A) and colony counts (CFU/mL) at CFS/2 and CFS/4 as compared to the untreated control. As the concentration of CFS decreased, colony counts become close to the untreated control (Figure 2B). The untreated groups generally exhibited high bacterial counts, ranging from approximately 10^8^ to 10^9^ CFU/mL. CFS treated condition resulted in varying degrees of bacterial reduction depending on the bacterial strain and CFS/MRS ratio. Overall, the antibacterial effect of CFS is strain-specific, with some pathogens showing substantial reductions in viable counts while others remained relatively higher in bacterial counts.

**Figure 2.**
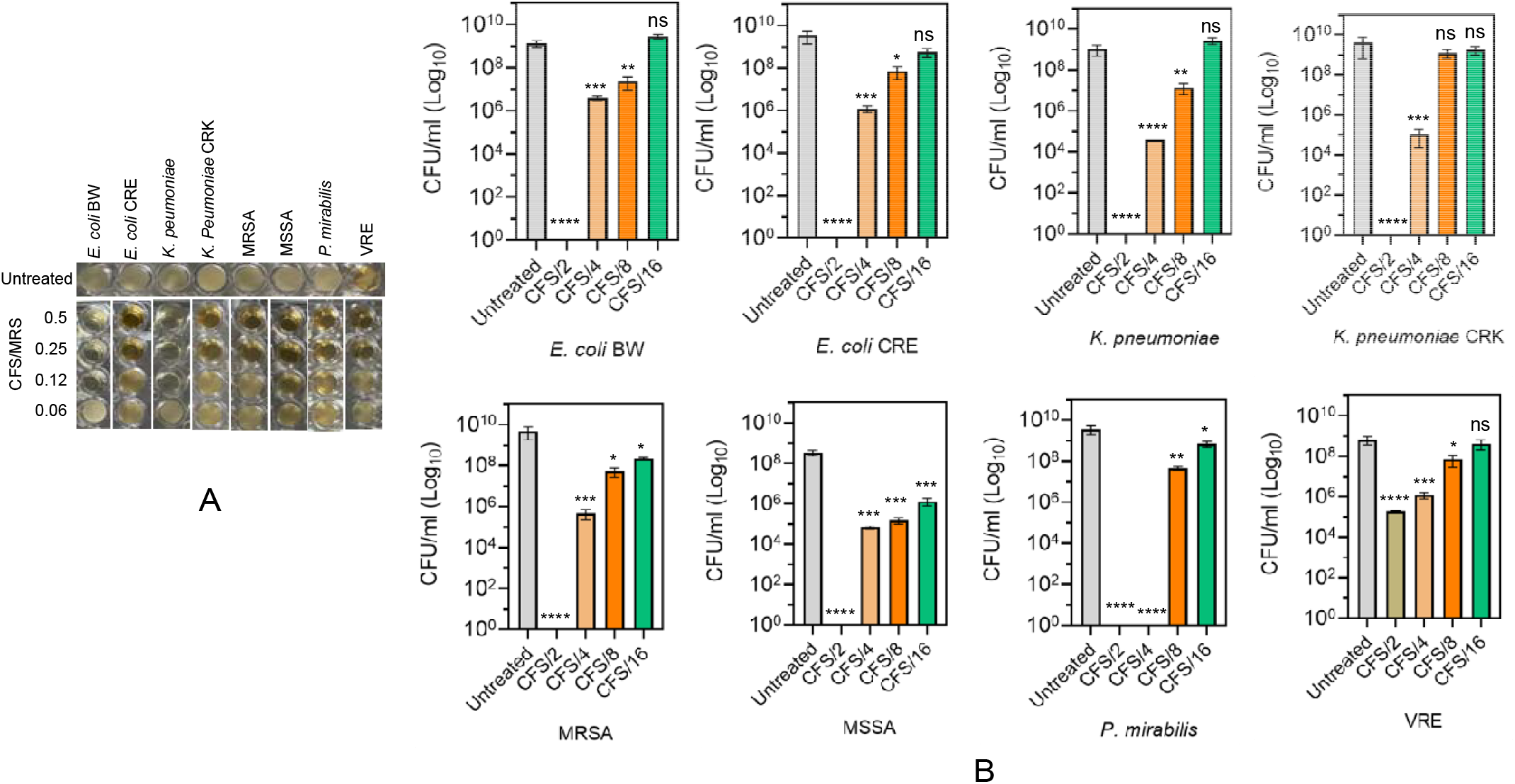
Impact of two-fold serial dilution on the antimicrobial potential of cell-free supernatant (CFS) against uropathogens. CFS was diluted in MRS broth to achieve final CFS/MRS v/v ratio of 0.5, 0.25, 0.12 and 0.06 (i.e. CFS/2, CFS/4, CFS/8 and CFS/16, respectively). 100 µL of Diluted CFS were mixed with 100 µL of different bacterial cell suspension in triplicate in 96-well microtiter plate and incubated for 24h at 150 RPM and 37□. Images of microtiter plate were taken using digital camera and cropped according to the bacterial lanes along with untreated control (A). Colony forming units (CFU) were determined after serial dilution for respective treatment conditions of each bacterium and represented as mean± SD of CFU/m on log10 scale (B).

The antibacterial effect of CFS of pH4 against all the selected eight uropathogen shows significant reduction in bacterial growth (Figure 3A) and CFU/mL (Figure 3AB) as compared to the untreated control. The CFS of pH5 also showed significant reduction in bacterial counts in comparison to untreated control. However, pH6 showed a slightly reduction in CFU of some uropathogens, while pH7 shows equivalent CFU/mL to untreated control (Figure 3A,B). These observations suggest that acidic pH is required to suppress the bacterial growth, and antimicrobial activity of CFS from *L. paracasei* SRG is pH dependent.

**Figure 3.**
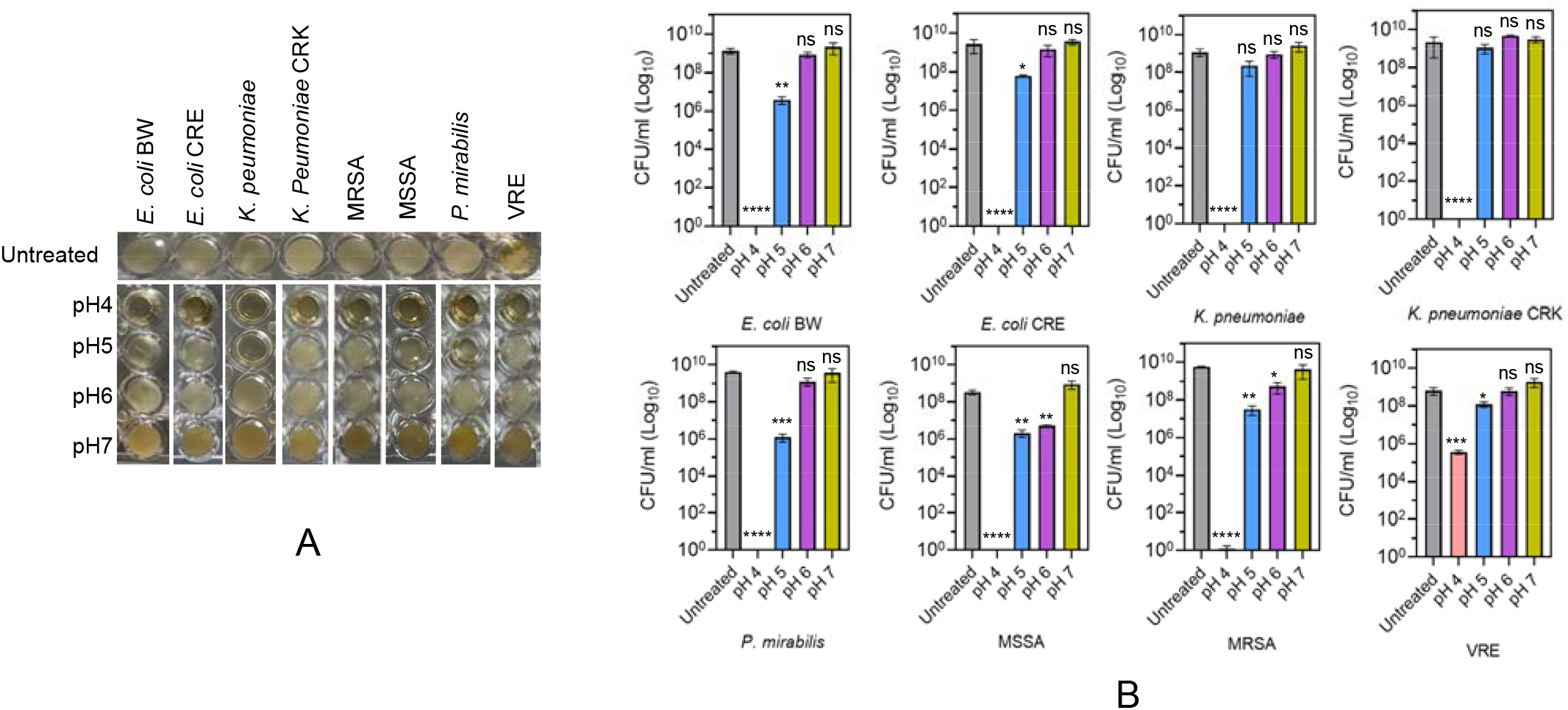
Impact of alkalinization on the antimicrobial potential of CFS against uropathogens. CFS/2 with pH4 was used for alkalinization, which was changed to pH5, pH6 and pH7 using 5N NaOH solution. 100 µL of CFS with different pH were mixed with 100 µL of different bacterial cell suspension in triplicate in 96-well microtiter plate and incubated for 24h at 150 RPM and 37□. Images of microtiter plate were taken using digital camera and cropped according to the bacterial lanes along with untreated control (A). Colony forming units (CFU) were determined after serial dilution for respective treatment pH conditions of each bacterium and represented as mean± SD of CFU/m on log10 scale (B).

To enumerate the role of bacteriocin in CFS of *L. paracasei* SRG, the antibacterial activity of CFS was monitored after heat inactivation and proteinase K enzymatic treatment. Under different treatment conditions viz. CFS, CFS + Pr K, and CFS + Heat, all the tested uropathogenic strains exhibited significant reduction in growth as visualized in microtiter plate image as well as colony counts (CFU/mL) as compared to untreated and MRS + Heat controls (Figure 4A,B). Importantly, the acidified MRS (pH4) has also showed significant reduction in bacterial counts in comparison to untreated control and heated MRS (Figure 4B). Altogether, these findings underscore that pH-dependent antimicrobial properties of CFS from vaginal isolate *L. paracasei* SRG are mediated by heat- and protease resistant metabolites.

**Figure 4.**
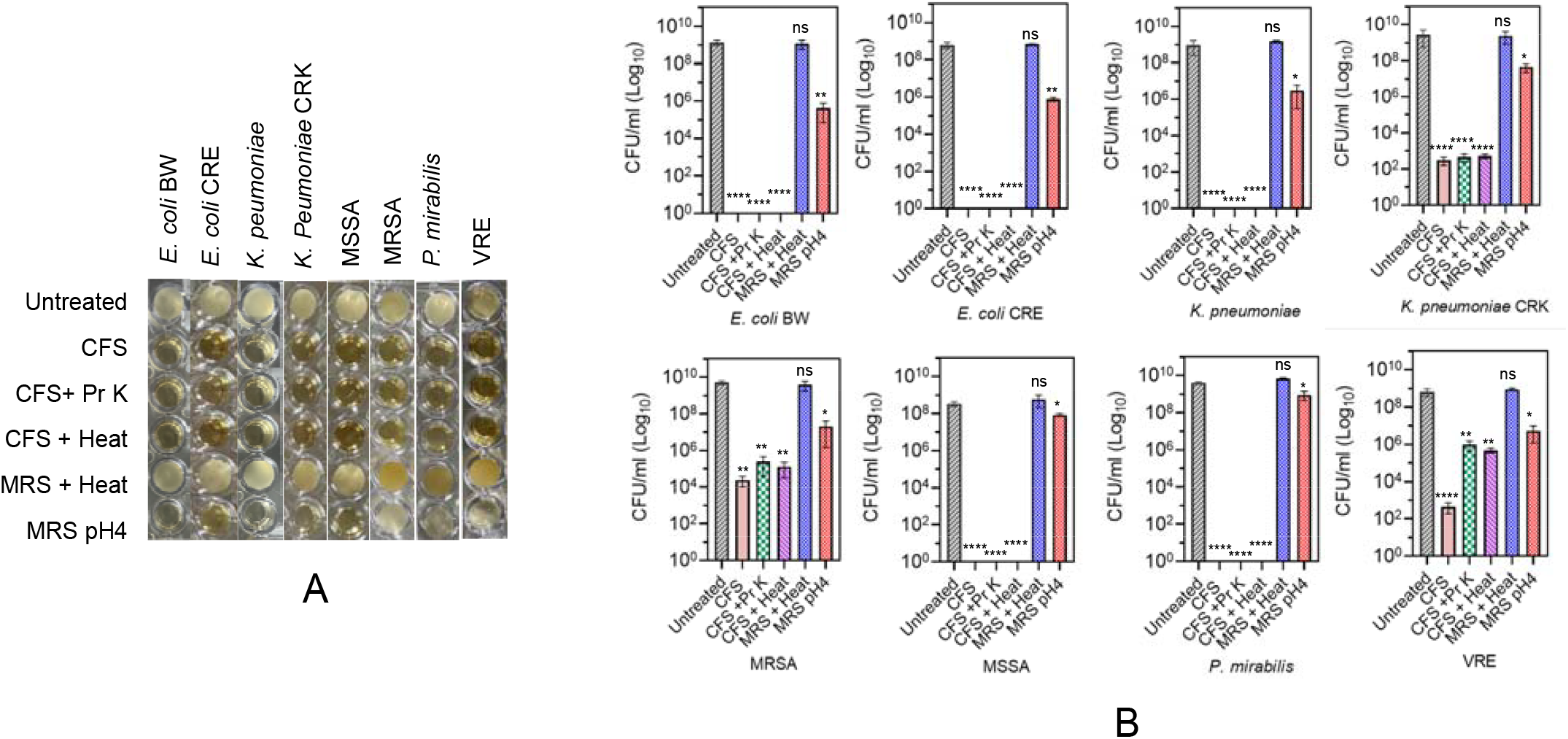
Effect of heat inactivation and protease treatment on antimicrobial activity of CFS from L. *paracasei* SRG. CFS was heat inactivated (98°C for 1h) or enzymatically treated with proteinase K (Pr K; 20 µg/mL for 1h at 37°C). Likewise, heated MRS broth and acidified MRS broth (pH4) was used as negative control. Next, 100 µL of treated CFS or MRS with 100 µL of different bacterial cell suspension in triplicate were incubated in 96-well microtiter plate for 24h at 150 RPM and 37□. Images of microtiter plate were taken using digital camera and cropped according to the bacterial lanes along with untreated control (A). Colony forming units (CFU) were determined after serial dilution for respective treatment conditions of each bacterium and represented as mean± SD of CFU/m on log10 scale (B).

### 3.4 Aggregation, co-aggregation and hydrophobicity

The autoaggregation and hydrophobicity percentages of *L. paracasei* SRG and the selected uropathogens are shown in Table 1. *L. paracasei* SRG showed the highest autoaggregation ability (85.32 ± 2.01%). Among the uropathogens, *S. aureus* JE2 demonstrated the highest autoaggregation (70.80 ± 3.02%), followed by *E. coli* CRE (65.60 ± 2.01%), *K. pneumoniae* CRK (64.80 ± 3.28%), and *K. pneumoniae* KCTC (62.99 ± 2.99%). In contrast, *P. mirabilis* MCC4207 showed the lowest autoaggregation among the tested pathogens (54.22 ± 4.55%). These findings indicate that autoaggregation varied among the bacterial strains, with *L. paracasei* SRG exhibiting substantially higher autoaggregation than the uropathogens examined. Higher autoaggregation is a probiotic feature, which may enhance the ability of *L. paracasei* SRG to form bacterial aggregates for better colonization and persistence at mucosal surfaces, and can promote biofilm formation and competitive exclusion of uropathogens from the vaginal and urinary tract mucosa [17,18]. However, these potential protective effects require further experimental confirmation through *in-vitro* and *in-vivo* assays.

**Table 1:** Autoaggregation and hydrophobicity percentages for *L. paracasei* SRG and pathogens.

| **Bacterial strains** | **Auto-aggregation (%)** | **Hydrophobicity (%)** |
| --- | --- | --- |
| *L. paracasei* SRG | 85.32 ± 2.01 | 55.6 ± 4.66 |
| *E. coli* BW25113 | 55.15 ± 1.98 | 29.33 ± 3.95 |
| *E. coli* CRE | 65.6 ± 2.01 | 34.11 ± 2.99 |
| *K. pneumoniae* KCTC 12385 | 62.99 ± 2.99 | 33.98 ± 3.85 |
| *K. pneumoniae* CRK | 64.8 ± 3.28 | 37.22 ± 5.25 |
| *P. mirabilis* MCC 4207 | 54.22 ± 4.55 | 5.66 ± 3.25 |
| *S. aureus* ATCC 25932 | 57.6 ± 1.98 | 49.33 ± 2.33 |
| *S. aureus* JE2 | 70.8 ± 3.02 | 54.33 ± 12.1 |
| *E. faecium* ATCC 700221 | 56.22 ± 3.15 | 18.99 ± 3.22 |

The cell surface hydrophobicity exhibited by *L. paracasei* SRG is 55.60 ± 4.66%. Among the uropathogens, *S. aureus* JE2 showed the highest hydrophobicity (54.33 ± 12.10%), followed by *S. aureus* ATCC25332 (49.33 ± 2.33%), *K. pneumoniae* CRK (37.22 ± 5.25%), *E. coli* CRE (34.11 ± 2.99%), *K. pneumoniae* KCTC (33.98 ± 3.85%), *E. coli* BW25113 (29.33 ± 3.95%), *E. faecium* ATCC70012 (18.99 ± 3.22%), and *P. mirabilis* MCC4207 showed the lowest value (5.66 ± 3.25%).

Co-aggregation percentage of *L. paracasei* SRG strain with pathogenic bacteria represented in Table 2. *L. paracasei* SRG exhibited co-aggregation with all eight selected uropathogens, with values ranging from 56.99 ± 5.66% to 71.62 ± 0.98%. The highest co-aggregation percentage was observed with *P. mirabilis* MCC4207 (71.62 ± 0.98%), followed by *K. pneumoniae* CRK (67.20 ± 3.01%), *S. aureus* JE2 (66.98 ± 7.85%), and *E. coli* CRE (66.36 ± 4.12%). Co-aggregation with *E. faecium* ATCC 70012 was 62.99 ± 6.21%, while *S. aureus* ATCC 25923 and *K. pneumoniae* KCTC exhibited values of 62.33 ± 5.99% and 60.21 ± 3.90%, respectively. The lowest co-aggregation was observed with *E. coli* BW25113 (56.99 ± 5.66%). Higher hydrophobicity and stronger co-aggregation potential of *L. paracasei* SRG with indicator strains may facilitate close interactions of vaginal isolate with uropathogens, potentially hindering their adhesion to uroepithelial cells and reducing pathogen colonization. Collectively, it may suggest the potential of *L. paracasei* SRG as a candidate probiotic for UTI prevention.

**Table 2:**
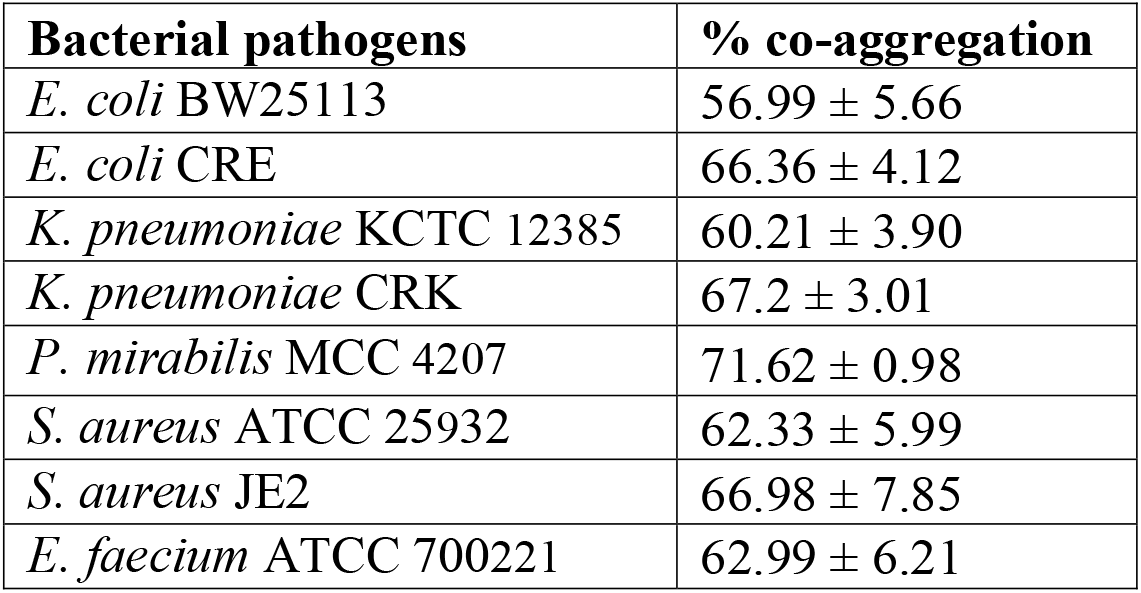
Co-aggregation percentage of *L. paracasei* SRG strain with pathogenic bacteria.

## 4. Conclusions

The vaginal isolate ‘*L. paracasei* SRG’ from healthy women and its CFS shows strong antimicrobial activities against several clinically important antibiotic resistant uropathogenic bacteria viz. *E. coli* BW25113, *E. coli* CRE, *K. pneumoniae* KCTC1285, *K. pneumoniae* CRK, *P. mirabilis* MCC4207, *E. faecium* ATCC700221, *S. aureus* JE2 and *S. aureus* ATCC 25923. The CFS from *L. paracasei* SRG exhibited strain-specific, pH-dependent antibacterial activity against uropathogens, primarily mediated by acidic conditions and heat- and protease-resistant metabolites. *L. paracasei* SRG showed strong autoaggregation ability (85.32 ± 2.01%), moderate cell surface hydrophobicity (55.60 ± 4.66%), and substantial co-aggregation with diverse bacterial pathogens. Overall, these findings support further investigation of *L. paracasei* SRG as a potential probiotic candidate developed from indigenous healthy vaginal microbiota, which may be used as an effective biotherapeutic to mitigate UTI and bacterial vaginosis. To validate its efficacy and potential applications, further, safety and *in vivo* studies are needed.

## Acknowledgments

R.A. gratefully acknowledges NBCFDC for NFOBC senior research fellowship. G.K.M. acknowledges IoE, BHU, UGC, India, and SERB-DST, India (SRG/2021/000900) for financial support. We acknowledge the contribution of Lipsa Behera during the sample collection and initial screening of isolates for their antimicrobial potential. We are thankful to Sukriti Chauhan and Rajnikant Roshan for their help in imaging of microtiter plate.

## Funding

This research received no external funding.

## Author’s contributions

R.A. conceptualized the study, performed the experiment, performed the analyses, prepared the figures, and wrote the main manuscript text. D.R. performed the experiment, contributed in analysis and manuscript writing. P.P. guided and reviewed the experiments. G.K.M. supervised the work, conceptualized the study, discussed the study, contributed in formal data analysis and final editing of manuscript. All authors read and approved the final manuscript.

## Competing interests

The authors declare no competing interests.

## Notes

### Competing Interest Statement

The authors have declared no competing interest.

